# PAC-MAP: Proximity Adjusted Centroid Mapping for Accurate Detection of Nuclei in Dense 3D Cell Systems

**DOI:** 10.1101/2024.07.18.602066

**Authors:** Tim Van De Looverbosch, Sarah De Beuckeleer, Frederik De Smet, Jan Sijbers, Winnok H. De Vos

## Abstract

**Motivation:** In the past decade, deep learning algorithms have surpassed the performance of many conventional image segmentation pipelines. Powerful models are now available for segmenting cells and nuclei in diverse 2D image types, but segmentation in 3D cell systems remains challenging due to the high cell density, the heterogenous resolution and contrast across the image volume, and the difficulty in generating reliable and sufficient ground truth data for model training. Reasoning that most image processing applications rely on nuclear segmentation but do not necessarily require an accurate delineation of their shapes, we implemented PAC-MAP, a 3D U-net based method that predicts the position of nuclei centroids and their proximity to other nuclei.

**Results:** We show that our model outperforms existing methods, predominantly by boosting recall, especially in conditions of high cell density. When trained from scratch PAC-MAP attained an average F1 score of 0.793 in dense spheroids. When pretraining using weakly supervised bulk data input and finetuning with few expert annotations the average F1 score could be significantly improved up to 0.817. We demonstrate the utility of our method for quantifying the cell content of spheroids and mapping the degree of glioblastoma multiforme infiltration in cerebral organoids.

**Availability and implementation:** The code is available on GitHub, at https://github.com/DeVosLab/PAC-MAP.

**Contact:** Winnok H. De Vos

## 1 Introduction

3D cell systems, such as explants, micro-tissues and organoids, are the next generation model systems for fundamental, preclinical and developmental research. Owing to their self-organizing nature, 3D cell systems can mimic part of the *in vivo* conditions, such as mechanical cues, chemical gradients, and patterning. Nonetheless their complexity demands more sophisticated and high-resolution methods for quality control and in-depth phenotypic characterization [1,2]. Light sheet fluorescence microscopy (LSFM) allows for acquiring images of intact samples at cellular resolution. By using cell type- and cell state-specific markers, the complete composition of such samples can in principle be visualized. However, automated analysis is hampered by the difficulty to automatically identify individual cells *in toto*. Conventionally, this challenge is tackled with cell segmentation based on image processing techniques that include thresholding, morphological operations, blob detection, edge detection, and seeded watershed or Voronoi segmentation to obtain cell masks [3]. However, deep learning (DL) has revolutionized cell segmentation. State-of-the-art methods, such as Stardist and Cellpose, provide an effective solution for segmenting nuclei or cells acquired with different microscopy modalities, and are well established for 2D images [4–6]. A shared feature of these methods is the prediction of spatial embeddings by a neural network, *e.g.,* gradients or vector fields, that allows for delineating individual cells via post-processing. These methods are based on supervised training and require the target spatial embeddings to be derived from manual annotations. While 2D annotated datasets are increasingly available [7,8], they are still scarce for 3D images due to the labor-intensive nature of accurate 3D image annotation. Moreover, because of the high cell density in 3D samples, even manually delineating individual cells is extremely challenging and requires specialized methods and sufficiently high resolution [9–11]. Therefore, DL-based 3D instance segmentation methods [12,13] are often limited to nuclei and tailored to specific datasets, and even here, annotation remains a major bottleneck. While in theory, instance positional information is less informative than instance segmentation, in practice, we argue, it is almost equivalent for dense 3D cell cultures, given 1) the difficulty in delineating cell bodies, 2) the prior knowledge that there is a considerable consistency in nuclei size, and 3) the notion that the direct region around the nuclei carries a significant part of phenotypic information on the cell [14–18].This raises the question as to whether instance position prediction could provide a more pragmatic, but equally valuable approach.

Recent works have investigated a DL based approach for nuclei detection using partial ground truth. As a compromise between retaining the accuracy of segmentation-based approaches and efficiency in data annotation, Krupa et al. [19] annotated the center 2D plane of nuclei and trained a 3D U-Net to segment the sparsely annotated foreground. Individual instances were then identified by thresholding the foreground probability and extracting the centroids. To further reduce data annotation efforts, Guo et al. [20] used point annotations for nuclei centroids to train a 3D U-Net to predict nuclei density maps. While both approaches outperformed conventional blob detection algorithms, they struggled with separating densely packed nuclei as they rely on connected component analysis after applying a fixed threshold.

To solve this, we replaced the nuclei density map with centroid probability maps and directly identified nuclei centroids as local maxima in the centroid probability map. Second, to better generalize to other datasets and to increase the resolution that can be achieved to separate nuclei, we train the neural network to jointly predict the centroid location and proximity to the nearest neighbor. Hereto, we adjust our targets by weighing the mean intensity of each Gaussian kernel to the distance to its nearest neighbor. The use of a weighted centroid probability map allows for an adaptive local maximum finding. We also tested whether cell detection performance can be improved by performing weakly supervised pretraining, using weak targets obtained from a conventional image processing algorithm, and subsequently finetuning on expert annotations. We demonstrate its value for cell counting in spheroids, identifying prelabeled subpopulations in co-culture spheroids and mapping the invasion of prelabeled glioblastoma multiforme cells in cerebral organoids.

## 2 Materials and methods

### 2.1 General procedure

A 3D U-Net model was trained to predict 3D nuclei centroid maps from 3D nuclear counterstain images (Fig. 1). Hereto, pairs of input images and target centroid maps were generated via manual point annotation of nuclei centroids. In inference mode, nuclei centroids were extracted by identifying local maxima in the predicted centroid maps. First, local maxima candidates were identified by finding the voxels that retained their value after performing a maximum filter with a spherical kernel with a radius equal to average radius of the nuclei in the dataset. Optionally, candidates with an intensity below a certain minimum value were discarded. From this stage forward, we make a distinction between the approaches that use a normal (C-MAP) and proximity-adjusted (PAC-MAP) probability map.

**Figure 1.**
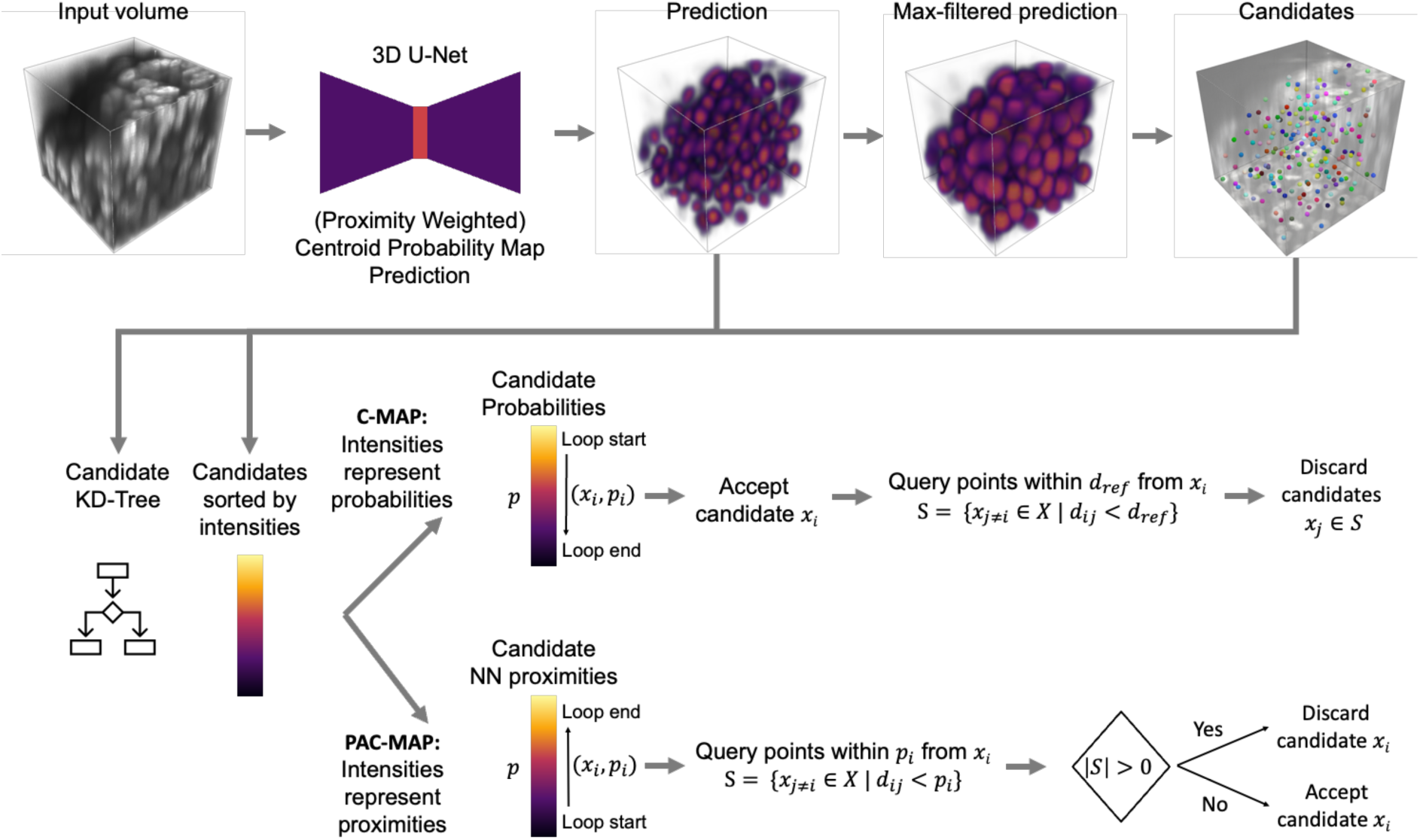
Nuclei detection via centroid probability prediction. First, A 3D U-Net-based model predicts a centroid probability map. Secondly, candidate centroids are identified as local maxima by performing a maximum filter and selecting the voxels that retained their values after filtering. Finally, final centroid predictions are obtained by filtering candidates in a for-loop based on their intensity and position relative to other candidates. Here we distinguish two appraoches: C-MAP and PAC-MAP. In C-MAP, the model predicts centroid probabilities, candidates are sorted in descending order based on their predicted probability. In each iteration, the top candidate is accepted and any candidates within a distance *d_ref_* are discarded. In PAC-MAP, the model predicts a nearest neighbor proximity adjusted centroid probability map, candidates are sorted in ascending order based on the predicted proximity to their nearest neighbor, and in each iteration, the top candidate is accepted only if no other candidates are found within its predicted proximity distance.

In C-MAP, the centroid map represents centroid probabilities and targets are created by positioning a Gaussian kernel on each annotation centroid position. Extracted centroid candidates are processed from high to low peak probability. To prevent multiple local maxima from being found within the same nucleus, a minimum reference distance *d_ref_* is defined to discard candidates closer than this distance from other candidates with higher peak probabilities. We defined *d_ref_* equal to the average radius of the nuclei in the dataset. Since setting this value for each dataset can be cumbersome and dependent on the cell density, we designed PAC-MAP as a more general alternative. In PAC-MAP, the centroid probabilities are adjusted by weighting each Gaussian kernel associated to a centroid with the proximity to the nearest neighbor. Hereby, the model learns to jointly predict centroid location and neighbor proximity. Extracted centroid candidates are processed from high to low predicted proximity and these proximities are used as local distance threshold. A candidate is discarded if other candidates are found within its associated predicted proximity threshold.

### 2.2 Generation of spheroids and organoids

Prior to spheroid and organoid production, cells were cultured following the procedures described in the Supplementary Information section. Spheroid and organoid production was performed in U-bottom 96 well plates (Sigma-Aldrich, CLS7007-24EA) coated with anti-adherence solution for 20 minutes (Stemcell technologies, 07010). The appropriate number of cells was seeded (between 2,000 and 10,000 cells/well) in 100 µl medium/well. Spheroids from SH-SY5Y or LN18-RED cells were left to form for at least three days before fixation. For cerebral organoid production, 10,000 NPCs were seeded. After seeding, the plates were centrifuged for three minutes at 100g before incubation at 37°C and 5 % CO_2_. Organoids were maintained for seven days (DIV7) with daily medium changes. Infiltration of GSC in organoids was achieved by co-culturing GSCs and cerebral organoids at DIV7. Prior to co-culturing with the organoids, the GSCs were dissociated using Tryple Express Enzyme and the desired GSC number was diluted in GSC medium. Medium was removed from the organoids and the GSCs were plated in GSC-medium onto the organoids. The GSCs were left to infiltrate the organoids for thirteen days before fixation. Sample fixation was performed by adding 1:1 volume 4 % PFA (ROTI-Histofix 4 % paraformaldehyde, Roth, 3105.2) overnight. Samples were washed three times with PBS after fixation and stored in PBS + 0.1 % NaN_3_.

### 2.3 Light sheet fluorescence microscopy image acquisition

Prior to imaging, all fixed samples (spheroids or organoids) stained with SiR-DNA (1.0 µM) nuclear counterstain (Spirochrome AG, SC007). Thereafter, the samples were embedded in 2% ultrapure agarose (Invitrogen, 16500500) using a custom-made cuboid mold. The resulting agarose cubes were placed in a glass-bottom dish and cleared overnight using the Fast 3D Clear protocol[21] (IBL Baustoff + Labor GmbH 220.110.021). The next day, the samples were mounted between the 5 mm water immersion TwinFlect mirrors (Leica, 158007011) of a Leica TCS SP8 Digital LightSheet microscope using a combination of a 2.5x illumination (506 523, fluostar, NA 0.07) and 10x detection objective (506 524 APO NA 0.30). We used a white light laser set at 488 nm, 552 nm, and 628 nm for excitation of GFP, RFP and SiR-DNA, respectively. Images were acquired using a voxel size (X, Y, Z) of 0.3594 ⨉ 0.3594 ⨉ 1.9999 µm^3^. An overview of all acquired LSFM datasets is presented in Table 1.

**Table 1.** Overview of all used datasets. LSFM: Light sheet fluorescence microscopy. For the average sample sizes of patches and spheroids, we list the patch size (X ⨉ Y ⨉ Z) and average measured diameter of the samples in the dataset, respectively.

| | Dataset | Type | Purpose | Source | # | Voxel size ($\mu\text{m}^3$) | Average sample size | Acquired Channels |
| --- | --- | --- | --- | --- | --- | --- | --- | --- |
| 1 | Mouse cortex training patches | LSFM | Training | [19] | 16 | $0.75 \times 0.75 \times 2.50$ | $84 \times 84 \times 80 \mu\text{m}^3$ | TO-PRO-3 |
| 2 | Mouse cortex evaluation patches | LSFM | Evaluation | [19] | 5 | $0.75 \times 0.75 \times 2.50$ | $150 \times 150 \times 150 \mu\text{m}^3$ | TO-PRO-3 |
| 3 | T47D training spheroids | Confocal | Training | [3] | 5 | $0.64 \times 0.64 \times 5.00$ | $D = 211 \pm 37 \mu\text{m}$ | SiR-DNA |
| 4 | T47D evaluation spheroids | Confocal | Evaluation | [3] | 3 | $0.64 \times 0.64 \times 5.00$ | $D = 174 \pm 5 \mu\text{m}$ | SiR-DNA |
| 5 | LN18-RED spheroids | LSFM | Pretraining | own | 6 | $0.3594 \times 0.3594 \times 1.9999$ | $D = 695 \pm 91 \mu\text{m}$ | SiR-DNA |
| 6 | SH-SY5Y spheroids | LSFM | Training | own | 2 | $0.3594 \times 0.3594 \times 1.9999$ | $D = 312 \pm 23 \mu\text{m}$ | SiR-DNA |
| 7 | Spheroids size range | LSFM | Proof-of-concept | own | 20 | $0.3594 \times 0.3594 \times 1.9999$ | $D = 929 \pm 310 \mu\text{m}$ | SiR-DNA |
| 8 | Co-culture spheroids | LSFM | Proof-of-concept | own | 8 | $0.3594 \times 0.3594 \times 1.9999$ | $D = 688 \pm 181 \mu\text{m}$ | eRFP<br>SiR-DNA |
| 9 | Organoid GSC infiltration | LSFM | Proof-of-concept | own | 2 | $0.3594 \times 0.3594 \times 1.9999$ | $D = 1077 \pm 272 \mu\text{m}$ | eGFP<br>SiR-DNA |

### 2.4 Data preprocessing

For all datasets, the intensities of the nuclear (SiR-DNA) channel were normalized and clipped to the range [0.0, 1.0] using the 0.1 and 99.9 percentiles, respectively. No normalization was performed on available RFP and EGFP marker channels. To reduce image size, images were cropped based on the nuclear channel using the bounding box around each sample. For our LSFM data, the images were subdivided in slightly overlapping patches of 256 ⨉ 256 ⨉ 46 voxels, or approximately 92 ⨉ 92 ⨉ 92 µm^3^ using a stride of 224, 224 and 40 voxels along the X-, Y-, and Z-axis. This patch size was chosen as a tradeoff between maximizing image and batch size and minimizing GPU memory usage during training.

### 2.5 Ground truth annotation

For model finetuning and evaluation, limited sets of respectively 30 and 18 images were manually annotated. Manual annotation was performed in Napari [22] by loading in a 3D image and creating a “points layer” to which annotated points, approximating the geometric center of the nucleus in X, Y and Z, were added. When using our method on the public datasets of Boutin et al. [3] and Krupa et al. [19], we used the geometric centroids of the segmented instances as ground truth. Since for the former dataset a training dataset was lacking, we additionally annotated five complete T47D (three days *in vitro*) spheroids.

### 2.6 Generation of target nuclei centroid probability maps

The target generation procedures are illustrated in Fig. 2. For C-MAP, centroid probability maps were generated from annotated centroids by positioning a 3D Gaussian kernel with an amplitude set to 1.0 and the standard deviation equal to the average nucleus radius encountered in the dataset onto the annotated centroids. The kernels were adjusted to the anisotropic voxel size of each dataset. For PAC-MAP, to establish a proximity-adjusted version of the probability map, pairwise distances between all centroids were first calculated, and the amplitudes of the Gaussian kernels were set equal to the distance to the nearest neighbor for each centroid.

**Figure 2.**
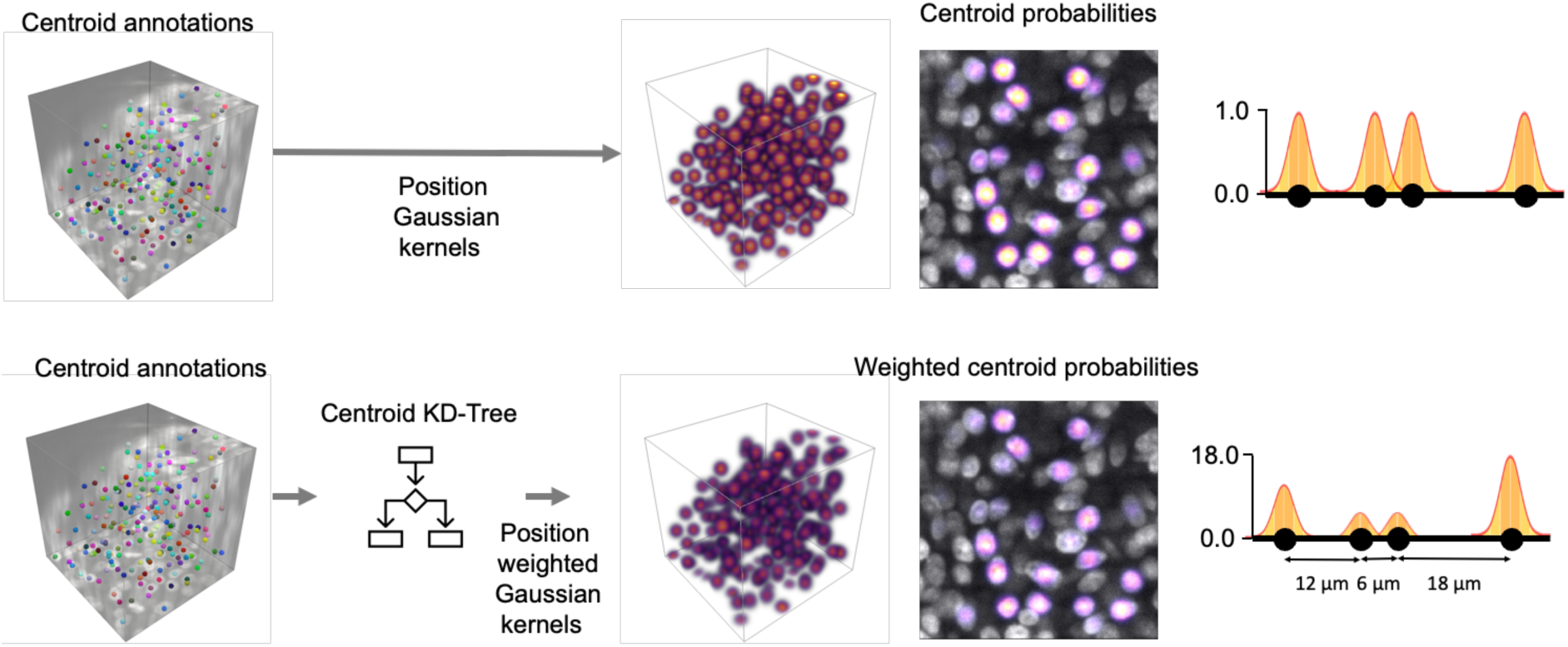
A: Target creation from annotated centroids. Top row: C-MAP - Centroid probability map creation by positioning Gaussian kernels at each centroid position. Bottom row: PAC-MAP - nearest neighbor proximity adjusted centroid probability map creation by positioning weighted Gaussian kernels at each centroid position. The weighted kernels are obtained by multiplying a Gaussian kernel with the distance to the nearest neighbor to each centroid.

### 2.7 Dataset generation

Three types of datasets with input-target pairs were generated: a pretraining dataset, a finetuning dataset and an evaluation dataset. For the pretraining dataset, estimates of nuclei centroids were obtained for all patches of the LN18-RED spheroids dataset (Table 1) by using a conventional image processing algorithm, including thresholding, distance map transformation, local intensity peak identification and seeded watershed segmentation. To avoid high background images from skewing the training dataset, patches with a foreground/background volume ratio below 1/3 were omitted from the pretraining dataset. For remaining patches (n = 2373), weak targets were obtained by positioning 3D Gaussian kernels at the centroids of the nuclei masks estimated by the seeded watershed-based algorithm. For creating the finetuning and evaluation datasets, respectively 30 and 18 patches were sampled from the SH-SY5Y spheroids dataset (Table 1). Nuclei centroids in these patches were manually annotated, and ground truth targets were generated by positioning (weighted) 3D Gaussian kernels at the annotated positions.

### 2.8 Model training

We used a 3D U-Net model [23] with a single input and output channel, an encoder depth of 4 layers with 32 feature maps for the initial first layer (doubling every layer), and a final sigmoid activation, as implemented by [24] in PyTorch. The MSE between target and predicted centroid probability maps was used as the loss function. In all cases, we used a batch size of 4, an initial learning rate of 1e^-4^, the Adam optimizer [25] with default parameters, and a learning rate scheduler that reduced the learning rate with a factor 0.5 if the loss had plateaued for 10 epochs.

Data augmentation was used in the form of random flipping on the X-, Y- and/or Z-axis, brightness adjustment with a mean of zero and a sigma of 0.1, contrast adjustment with a factor uniformly sampled between [0.9, 1.1], and Gaussian noise with a mean of zeros and a sigma of 0.1. All augmentation steps had a probability of 50%. We did not include random (anisotropic) rescaling, as it did not improve performance. To evaluate the reproducibility of model performance, all model training procedures were repeated three times with a different random seed (guaranteeing different but reproducible dataset compositions, model initializations, batch shuffling, and data augmentation transformations).

For the public datasets, we trained models from scratch on manual annotations (in case of [3]), or on the available training data (in case of [19]) (see section 2.5). For our own LSFM dataset, three types of models were generated: pretrained models trained with weak annotations; finetuned models resulting from finetuning the pretrained models on manual annotations; models trained from scratch on manual annotations. First, weakly supervised pretraining was done on the pretraining dataset for 100 epochs. The dataset was split at random in training and validation subsets with respectively 80 % and 20 % of the data. The model state with the lowest loss score on the validation set was selected and evaluated on the test data of manual annotated samples. Next, the pretrained models were finetuned for 50 epochs on the finetuning dataset using an 80% and 20% training and validation dataset split. The finetuned model states with the lowest loss scores on the validation set was then evaluated on the same evaluation dataset. Finally, we also trained models from scratch for 150 epochs on the finetuning dataset using an 80% and 20% training and validation dataset split and tested it on the evaluation dataset.

### 2.9 Evaluation

We compared predicted and ground truth centroids by first matching them via linear sum assignment [26], *i.e.,* the Hungarian algorithm, and calculated TP, FP and FN between matched predicted and ground truth points. Only matched predicted centroids within a distance of 5 µm were counted as TP. Unmatched predicted and ground truth centroids were counted as FP and FN, respectively. Likewise, matched predicted and ground truth centroids with a distance ≥ 5 µm, were also counted as FP and FN, respectively. Based on these scores, the precision, recall and F1-score were calculated (see equations 1, 2, and 3). We reported the average and standard deviation of model performance scores for each model training procedures. In case ground truth masks were available, we used the geometric centroid of each mask as the ground truth centroid.

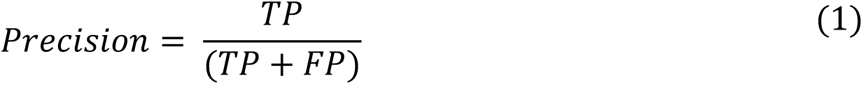

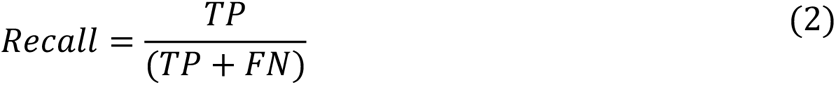

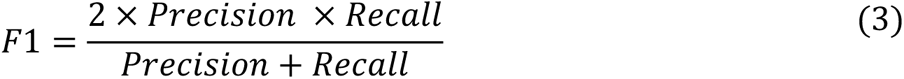

### 2.10 dentification of cell phenotypes and their radial position

After centroid prediction was performed based on the nuclear stain, cell phenotypes were identified using the available marker channel. Hereto, a threshold on the local average intensity of a marker within a sphere (r = 5 µm) centered at the predicted centroid was used. To identify the LN18-RED cells in LN18-RED/NPC co-culture spheroids, we used a fixed threshold on the average marker intensity. For each co-culture spheroid, we reported the resulting cell type counts and their ratio. For identifying GSCs in cerebral organoids, a fixed intensity threshold for EGFP did not work since we found a linear relationship between the intensity of the nuclear stain and the marker channel for non-GSCs cells. We therefore fitted a linear regression to this relation based on 10,000 randomly sampled NPC nuclei and applied a threshold at the upper bound of the 99.9 % confidence interval of this fit, instead. We visualized GSC invasion patterns in each organoid by quantifying the GCS/NPC ratio for each GSC in a 25 µm radius local neighborhood and plotting this ratio as a function of the radial position within the organoid. The radial position of each cell was obtained binarizing the foreground and extracting the value of the distance transform at the centroid location (*i.e.,* the distance to the surface).

### 2.11 Implementation details

All image preprocessing, model training and evaluation was implemented in Python 3.9 using numpy (1.24.3), pandas (2.1.0), scikit-image (0.21.0), scikit-learn (1.3.0), and PyTorch (2.0.1). To limit conflicts between packages, a separate conda environment was created for running Stardist (0.8.3). Model training and inference were performed on a Linux server system (Ubuntu 20.04.6 LTS) with an AMD EPYC 7282 16-Core processor using an NVIDIA RTX A6000 GPU.

## 3 Results

### 3.1 Proximity-weighted centroid probability maps are effective for nuclei detection

We hypothesized that cell detection in images of dense 3D cell cultures could be facilitated by predicting the location of nuclear centroids using a criterion that considers the local density of these points. We first tested our hypothesis on the publicly available LSFM mouse brain LSFM images stained with TO-PRO-3 [19]. The average density of this sample was 138.000 nuclei/mm^3^. We compared our method to two state-of-the-art methods (NuMorph, foreground prediction + connected components analysis [19], and SAU-Net, density map prediction + connected components analysis [20]). Note that Guo et al. [20] propose the addition of an attention module to the 3D U-Net architecture, but for simplicity and fair conceptual comparison between these methods, we used the same 3D U-Net backbone (4 levels, 32 starting feature maps) for all methods. It is reasonable to assume that improvements to the model architecture would benefit each method similarly. We trained all models from scratch on the available training data with the same hyperparameters and data augmentation strategies. For each method, model training was repeated trice with random assignment of the 16 training samples into a training and validation dataset (80 – 20 % split). Evaluation was performed on 5 unseen test samples. In addition, we compared to 3D Stardist [12] for which we used the publicly available demo model and used the geometric centroids of the segmented instances as the predicted centroids. For all methods, hyperparameters were set by optimizing the F1 score on the training set. Table 2 summarizes the performance of all methods on the test dataset. In terms of F1, our methods (C-MAP and PAC-MAP) outperformed all other evaluated methods. Note that NuMorph, SAU-Net and C-MAP rely on setting a threshold (either on intensity or distance between centroid candidates), which is not required for PAC-MAC that achieved either the highest or second-best score for all metrics. Only in terms of precision, SAU-Net scored the best at the cost of a higher FN rate. For a slight drop in precision, a centroid prediction-based method can thus significantly improve recall. The drop in precision is mostly caused by the false detection of multiple local maxima within a single nucleus, which is unlikely when connected components are extracted after thresholding. On the other hand, the latter has a higher risk of considering densely packed nuclei as one as they will stay connected after thresholding [19,20]. In terms of F1, the 3D Stardist model obtained the lowest performance, but it must be noted that this was a publicly available demo model that trained on simulated data. Fig. 3 visualizes a typical case from the test set in which touching nuclei resulted in undetected nuclei by NuMorph and SAU-Net, while C-MAP and PAC-MAP could resolve all touching nuclei. Thresholding of model predictions, a shared strategy of NuMorph and SAU-Net, resulted in merging of neighboring nuclei.

**Figure 3.**
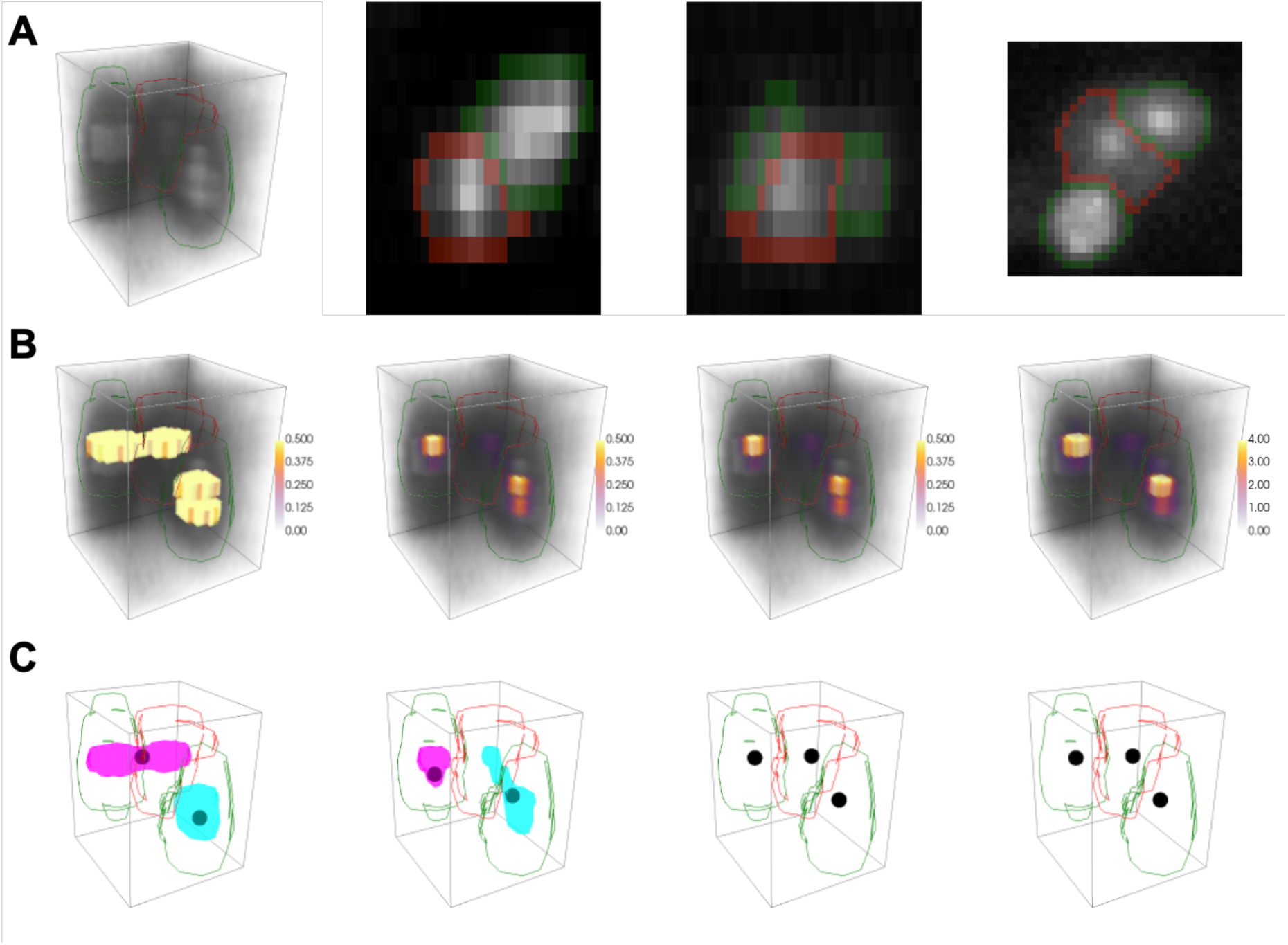
Visualization of advantage of nuclei centroid prediction compared to thresholding and connected component analysis. Images show a region with three touching nuclei in the mouse brain LSFM images [19]. A: Image volume (22.5 × 22.5 × 30.0 *μm*^3^) and orthogonal slices (XZ, YZ, and XY) with annotated nuclei (left to right). The two reen nuclei are detected by all methods, while the red nucleus is only detected using C-MAP and PAC-MAP; B: Model predictions by NuMorph, SAU-Net, C-MAP and PAC-MAP (left to right). For visualization purposes, intensities are clipped between 0 and 50% of the maximum value in the prediction volume; C: Connected components and/or predicted centroids extracted from model predictions in B for NuMorph, SAU-Net, C-MAP and PAC-MAP.

**Table 2.** Evaluation of nuclei centroid prediction on LSFM mouse brain data [19]. NuMorph: binary segmentation and connected components analysis (based on [19]); SAU-Net: nuclei density map prediction and connected components analysis (based on [20]); C-MAP: nuclei centroid prediction via probability map prediction; PAC-MAP: nuclei centroid prediction via proximity adjusted probability map prediction.

| Method | Precision ($\mu \pm \sigma$) | Recall ($\mu \pm \sigma$) | F1 ($\mu \pm \sigma$) |
| --- | --- | --- | --- |
| 3D Stardist | 0.974 | 0.878 | 0.924 |
| NuMorph | $0.968 \pm 0.007$ | $0.894 \pm 0.020$ | $0.929 \pm 0.014$ |
| SAU-Net | **$0.986 \pm 0.002$** | $0.917 \pm 0.004$ | $0.950 \pm 0.003$ |
| C-MAP | $0.978 \pm 0.002$ | **$0.946 \pm 0.003$** | $0.961 \pm 0.002$ |
| PAC-MAP | $0.979 \pm 0.006$ | $0.944 \pm 0.008$ | **$0.962 \pm 0.000$** |

### 3.2 Nuclei centroid prediction becomes more advantageous in dense samples

In spheroids and cerebral organoids, we observed a significantly higher average cell density of around 232.000 nuclei/mm^3^ compared to the mouse brain LSFM dataset. The higher density in such samples results in images in which nuclei appear to be touching and overlapping more often, which increases the difficulty of nuclei detection. We trained and tested the same detection methods on an annotated LSFM dataset of spheroids, except for NuMorph as manual ground truth nuclei segmentations were not available. The cell detection models outperformed the pretrained 3D Stardist model. PAC-MAP (F1 = 0.793 ± 0.011) outperformed the SAU-Net approach (F1 = 0.750 ± 0.039), again owing to a strong gain in recall (see Table 3). PAC-MAP also outperformed C-MAP for both precision and F1. Although overall performance dropped in these more challenging samples, the relative performance of PAC-MAP to SAU-Net, measured in terms of F1, increased from +1.3 % to +5.7 %. This shows the advantage of direct centroid identification via centroid probability prediction compared to using segmentation via thresholding in dense samples.

**Table 3.** Evaluation of nuclei centroid prediction on LSFM spheroid data. SAU-Net: nuclei density map prediction and connected components analysis (based on [20]); C-MAP: nuclei centroid prediction via probability map prediction; PAC-MAP: nuclei centroid prediction via proximity adjusted probability map prediction.

| Method | Precision ($\mu \pm \sigma$) | Recall ($\mu \pm \sigma$) | F1 ($\mu \pm \sigma$) |
| --- | --- | --- | --- |
| 3D Stardist | 0.650 | 0.687 | 0.658 |
| SAU-Net | **$0.808 \pm 0.005$** | $0.740 \pm 0.118$ | $0.750 \pm 0.039$ |
| C-MAP | $0.724 \pm 0.021$ | **$0.865 \pm 0.007$** | $0.783 \pm 0.014$ |
| PAC-MAP | $0.755 \pm 0.005$ | $0.847 \pm 0.017$ | **$0.793 \pm 0.007$** |

### 3.3 Weak supervised pretraining improves nuclear detection performance

While point annotations can be done much faster than manual nuclei segmentation, it is still labor intensive. Therefore, we tested if performance can be improved by pretraining a model weak targets produced by a conventional segmentation algorithm, which is assumed to be readily available for most datasets. Specifically, we used thresholding and seeded watershed segmentation to produce weak targets for pretraining (Fig. 4). For all methods, models were first pretrained for 100 epochs and then finetuned for 50 epochs on the annotated training dataset. In finetuning, we initiated the model with the weights of the checkpoint that had the best performance on the validation dataset of the pretraining task. Evaluating the seeded watershed-based predictions itself, we found that it resulted in missed centroids and that it also returned many false positive centroids in background regions due to segmentation errors. Interestingly, the model that was pretrained on the weak targets displayed superior performance as compared to this benchmark. Fig. 5 shows that the pretrained models detected centroids that were missed by the method by which it was supervised and that it returned less false positives. Quantitatively, all pretrained cell detection models (F1 > 0.584) performed significantly better than the seeded watershed-based predictions (F1 = 0.516) but were outperformed by the pretrained 3D Stardist model (F1 = 0.658) (see Table 4). As expected, all pretrained models obtained lower performance compared to models trained directly from scratch on 30 manually annotated patches (F1 > 0.750, see Table 3). However, the combination of pretraining and finetuning resulted in a significant improvement compared to training from scratch for all metrics (F1 ≥ 0.785). Both in terms of F1 and recall our approaches achieved the best performance (F1 ≥ 0.817, Recall ≥ 0.850). C-MAP and PAC-MAP achieved similar performance in F1, while outperforming the others in recall and precision, respectively. SAU-Net again obtained best precision (0.827 ± 0.012) at the cost of a lower recall (0.767 ± 0.013). In terms of average F1, the performance gains from finetuning over training from scratch for SAU-Net, C-MAP and PAC-MAP were respectively 4.7, 4.5 and 3.0 %. From the results above, we deduce that weak supervision, i.e., bootstrapping model training on inaccurate targets from a conventional image processing algorithm, could be a general strategy for pretraining prior to finetuning the models on a limited set of annotated samples.

**Figure 4.**
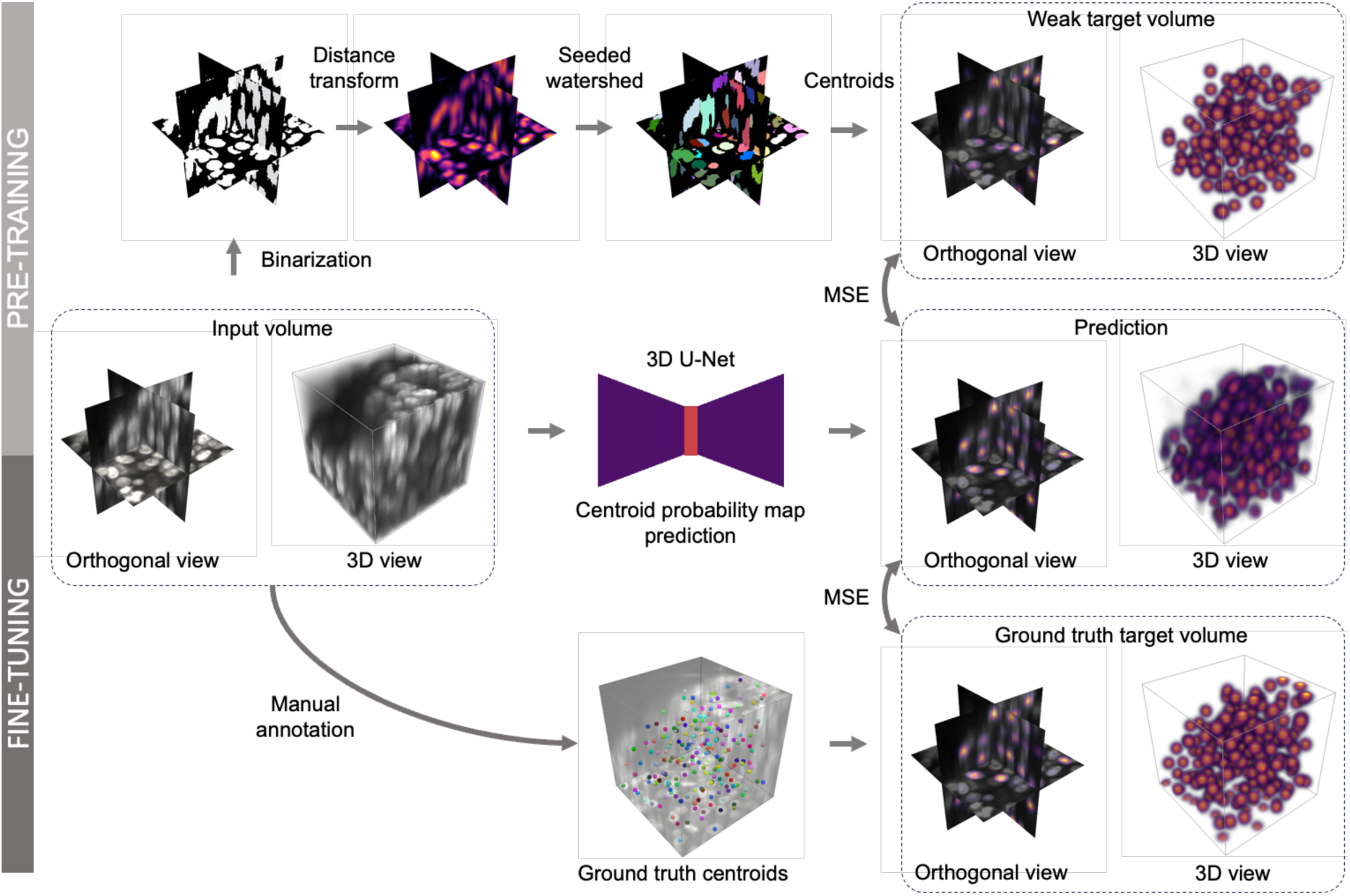
Model pretraining and finetuning workflow: 3D volumes of LN-18 and SH-SY5Y spheroids were subdivided into patches. Then, weakly supervised pretraining is used using the LN-18 dataset with (suboptimal) centroid estimations resulting from a conventional image processing algorithm. Finally, the model is finetuned using human annotations on the SH-SY5Y dataset.

**Figure 5.**
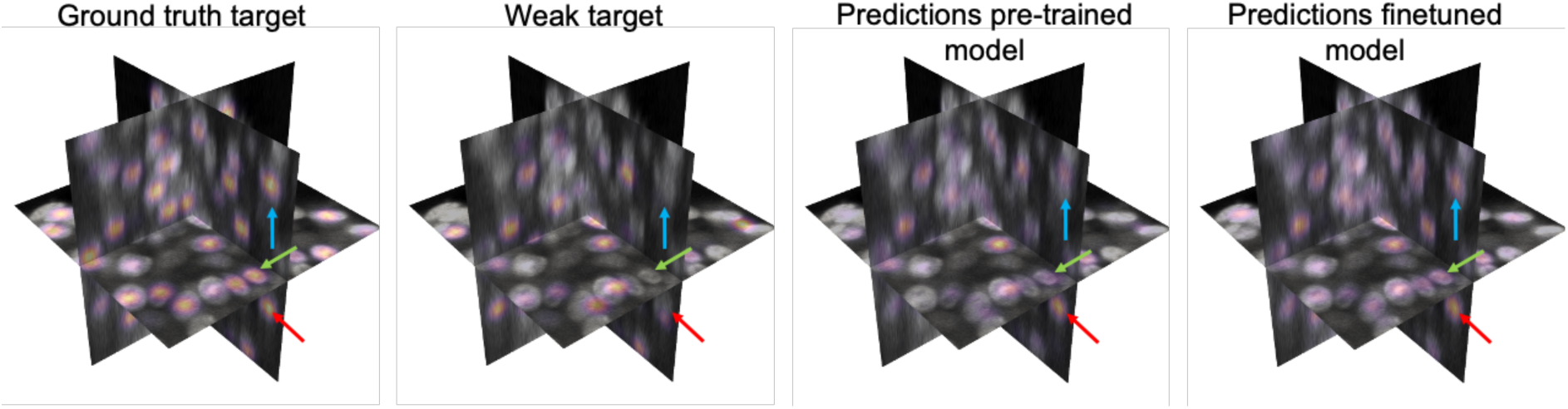
Orthogonal slices through the volume of an SH-SY5Y patch overlayed with the centroid probability maps obtained from ground truth annotation, the weak targets, the pretrained model predictions, and the finetuned model predictions. Arrows indicate nuclei undetected by the conventional image processing algorithm but detected by the pretrained and in greater extend by the finetuned model.

**Table 4.** Evaluation of pretrained and finetuned models in nuclei centroid prediction on LSFM spheroid data. SAU-Net: nuclei density map prediction and connected components analysis (based on [20]); C-MAP: nuclei centroid prediction via probability map prediction; PAC-MAP: nuclei centroid prediction via proximity adjusted probability map prediction.

| Method | Precision ($\mu \pm \sigma$) | Recall ($\mu \pm \sigma$) | F1 ($\mu \pm \sigma$) |
| --- | --- | --- | --- |
| Seeded watershed segmentation | 0.515 | 0.542 | 0.516 |
| 3D Stardist | 0.650 | 0.687 | 0.658 |
| SAU-Net pretrained | $0.600 \pm 0.028$ | $0.587 \pm 0.002$ | $0.584 \pm 0.013$ |
| C-MAP pretrained | $0.557 \pm 0.017$ | $0.671 \pm 0.002$ | $0.599 \pm 0.012$ |
| PAC-MAP pretrained | $0.550 \pm 0.004$ | $0.656 \pm 0.008$ | $0.590 \pm 0.003$ |
| SAU-Net finetuned | **$0.827 \pm 0.012$** | $0.767 \pm 0.013$ | $0.785 \pm 0.011$ |
| C-MAP finetuned | $0.781 \pm 0.009$ | **$0.876 \pm 0.004$** | **$0.818 \pm 0.004$** |
| PAC-MAP finetuned | $0.795 \pm 0.003$ | $0.850 \pm 0.010$ | $0.817 \pm 0.006$ |

### 3.4 Nuclei centroid prediction enables absolute quantification of spheroid cell content

As one of the main challenges in organoid research is reproducibility, sample composition analysis is crucial from a quality control perspective. Hence, to demonstrate the utility of direct nuclei centroid prediction, we used the optimal PAC-MAP model (in terms of F1 score on the evaluation dataset) to count all cells in spheroids seeded at different densities. For this purpose, a set of twenty light sheet image stacks of LN18-RED spheroids were acquired 72h after seeding 2000 (n = 2), 4000 (n = 3), 6000 (n = 7) and 8000 (n = 8). Automated cell detection retrieved correspondingly increasing cell numbers in the same order of magnitude but with an apparent blunting at the highest density (Fig 6A). Next, we analyzed the composition of eight co-culture spheroids that contained NPCs and LN18-RED cells, seeded at a 10:1 ratio and cultured for 72h, by quantifying the number of the two constituent cell types. We detected on average a much higher number of LN18-RED nuclei (8880 ± 1174) leading to a ratio of 0.67 ± 0.11 (Fig 6B), plausibly due their much faster growth rate. Thus, PAC-MAP enables absolute quantification of cell composition of dense 3D cell systems.

**Figure 6.**
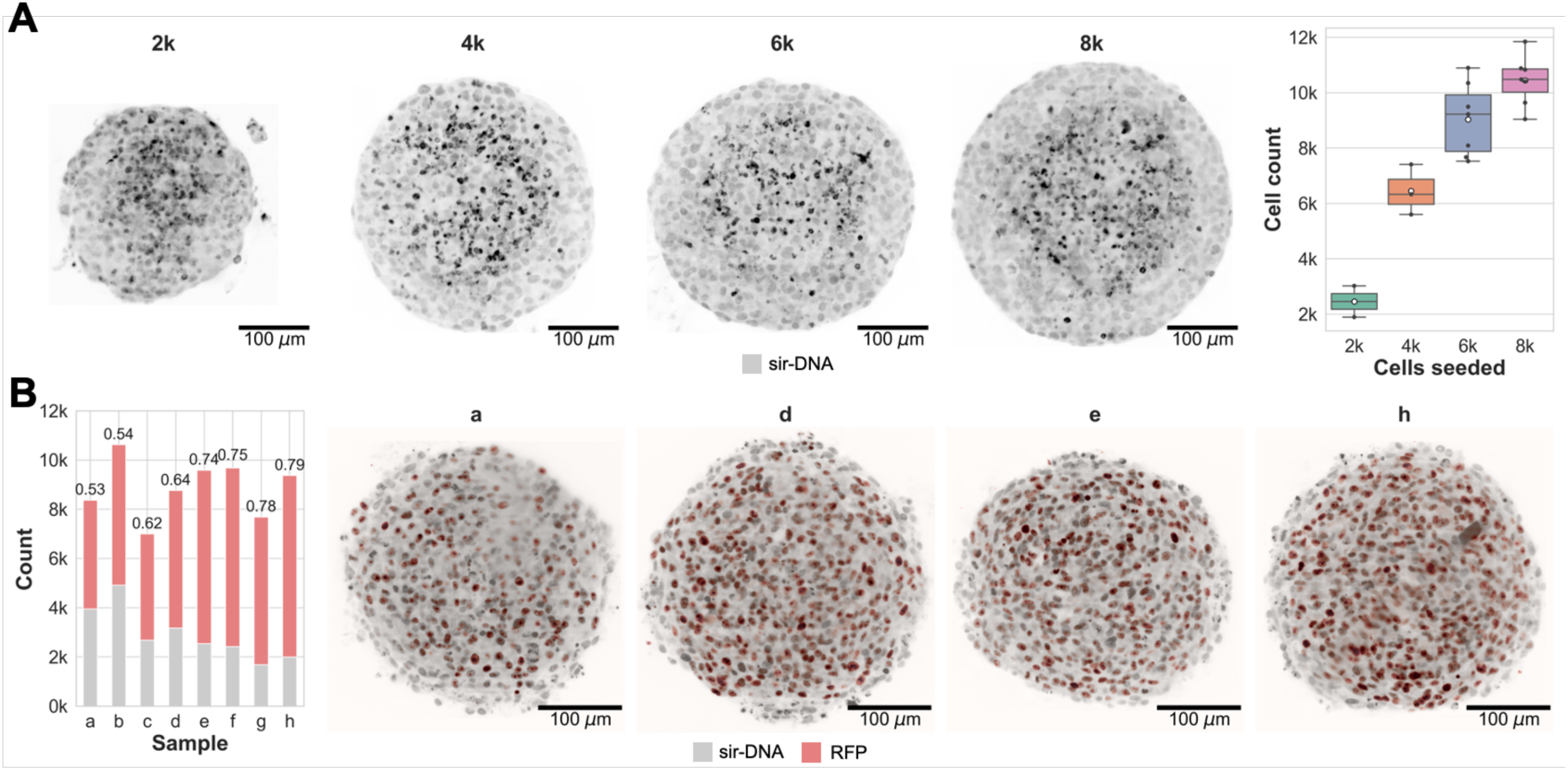
Quality control applications for spheroids. A: Measured cell count as function of number of seeded cells for LN-18-RED spheroids; B: Measured cell count and cell type ratio for spheroids created by seeding 10000:1000 LN-18-RED and NPC cells. LN-18-RED cells were identified based on their marker intensity (RFP).

### 3.5 Nuclei centroid prediction identifies changes following drug treatment

Using PAC-MAP, we next reproduced the results from [9] in which the number and radial position of mitotic cells in T47D spheroids were detected following nocodazole drug treatment at different concentrations using a nuclei segmentation-based approach. Mitotic cells were identified using the mitotic marker phospho-histone H3 (PH3). Since their images were acquired using a confocal microscope at a different voxel size (0.64 ξ 0.64 ξ 5 µm^3^), we first trained a nuclei centroid prediction model from scratch on their data. Hereto, we annotated five small randomly selected 3DIV T47D spheroids for training that were not part of the drug treatment experiment. We trained a smaller model (16 instead of 32 starting feature maps) for 150 epochs with a batch size of 12. As before, the annotated spheroids were subdivided in patches of which 80 and 20 % were used for model training and validation, respectively. After training, we evaluated the performance of our method in nuclei centroid prediction and compared it to their segmentation-based approach on three samples that were manually segmented by the original authors. Based on their evaluation criteria, our method (F1 = 0.83 ± 0.05) outperformed their segmentation-based approach (0.76). While both methods obtained similar precision of respectively 0.87 and 0.88, they differed substantially in recall (0.80 and 0.67). Next, we detected all nuclei in all samples part of the drug screen and measured to average PH3 intensity in a 5 µm radius around each nucleus. Using the same thresholds on average PH3 intensity and radial position as used by [9], the cell phenotypes (PH3+/-) and their layer (inner/outer) were identified. Without the need for instance segmentation, we could reproduce the distribution of PH3+ cells as a function of the spheroid layer and applied nocodazole concentration (Fig. 7).

**Figure 7.**
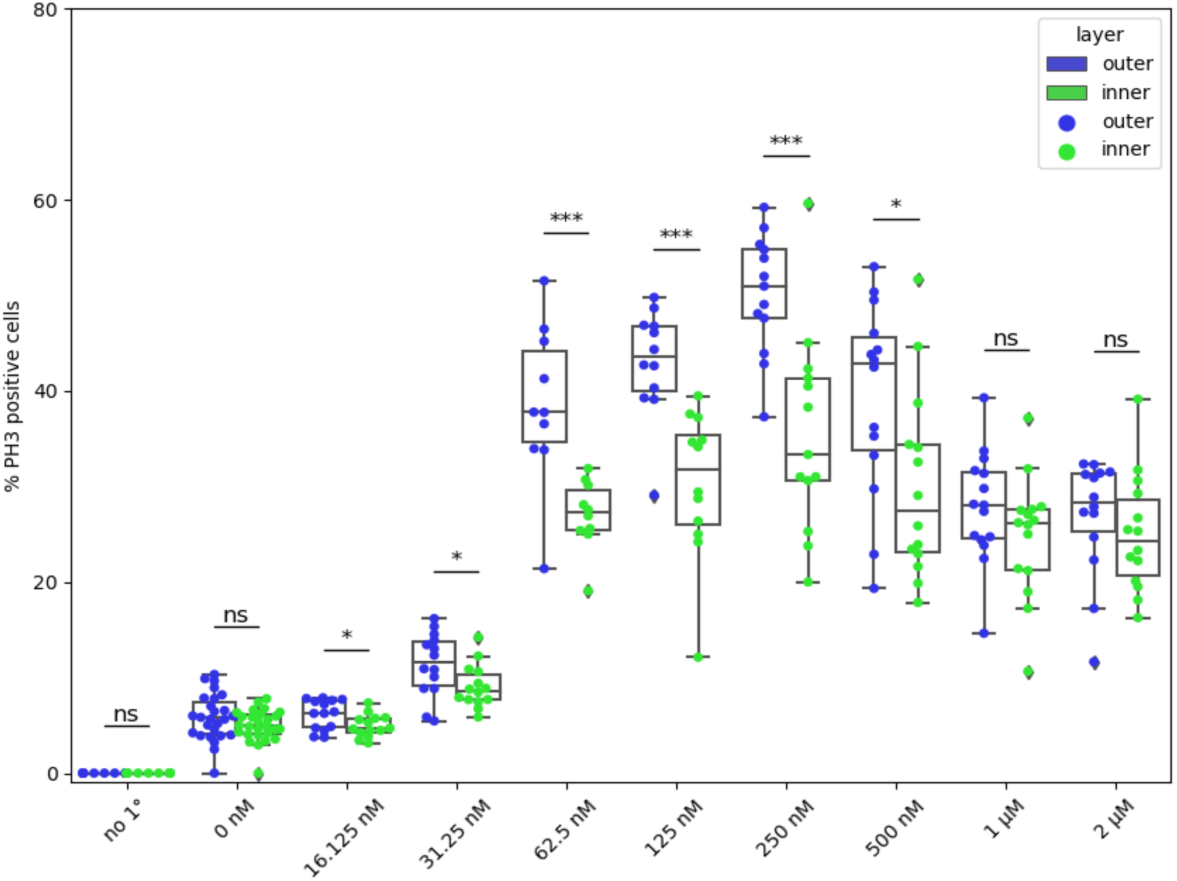
Reproduction of analysis by [3]. Measurement of subpopulations (PH3^+/-^ cells) within spheroids after treatment as a function of nocodazole treatment and radial layer. Eventhough we relied on centroid prediction only, our measurements closely resemble the onces published by [3] that were obtained using instance segmentation.

### 3.6 Nuclei centroid prediction allows mapping GSC infiltration in cerebral organoids

As a final application, we applied our approach to a biologically relevant use case with high preclinical relevance, *i.e.,* the mapping of glioma stem cell (GSC) location in cerebral organoids as a model for glioblastoma infiltration. Two cerebral organoids were seeded with respectively 1000 and 2000 EGFP-expressing GSC and imaged thirteen days after. At the time of fixation, the organoids had a diameter of around 1100 µm. After nuclei centroid prediction, the radial positions of all cells were obtained and GSCs were identified based on a threshold on the average GSC marker intensity (EGFP) within a sphere of 5 µm radius centered at the centroid. In the organoids seeded with 1000 and 2000 GSC cells, 1366 and 2587 GSCs where detected, respectively. For all GSCs, we performed a cell type composition analysis within the local 25 µm radius neighborhood in relation to their radial position. Since the samples are not perfectly spherical, we expressed radial position as distance to the surface (Fig. 8A). This analysis showed different GCS distributions between both samples (Fig. 8B and C). In the first sample, densely packed GSCs were found either close to the surface (<100 µm from the surface), or to the center. In the second sample, densely packed GSCs were found in between the center and the surface of the organoid. The second sample also contained more GSCs at its surface. For both samples, most isolated GSCs were found dispersed between 25 and 200 µm from their surface.

**Figure 8.**
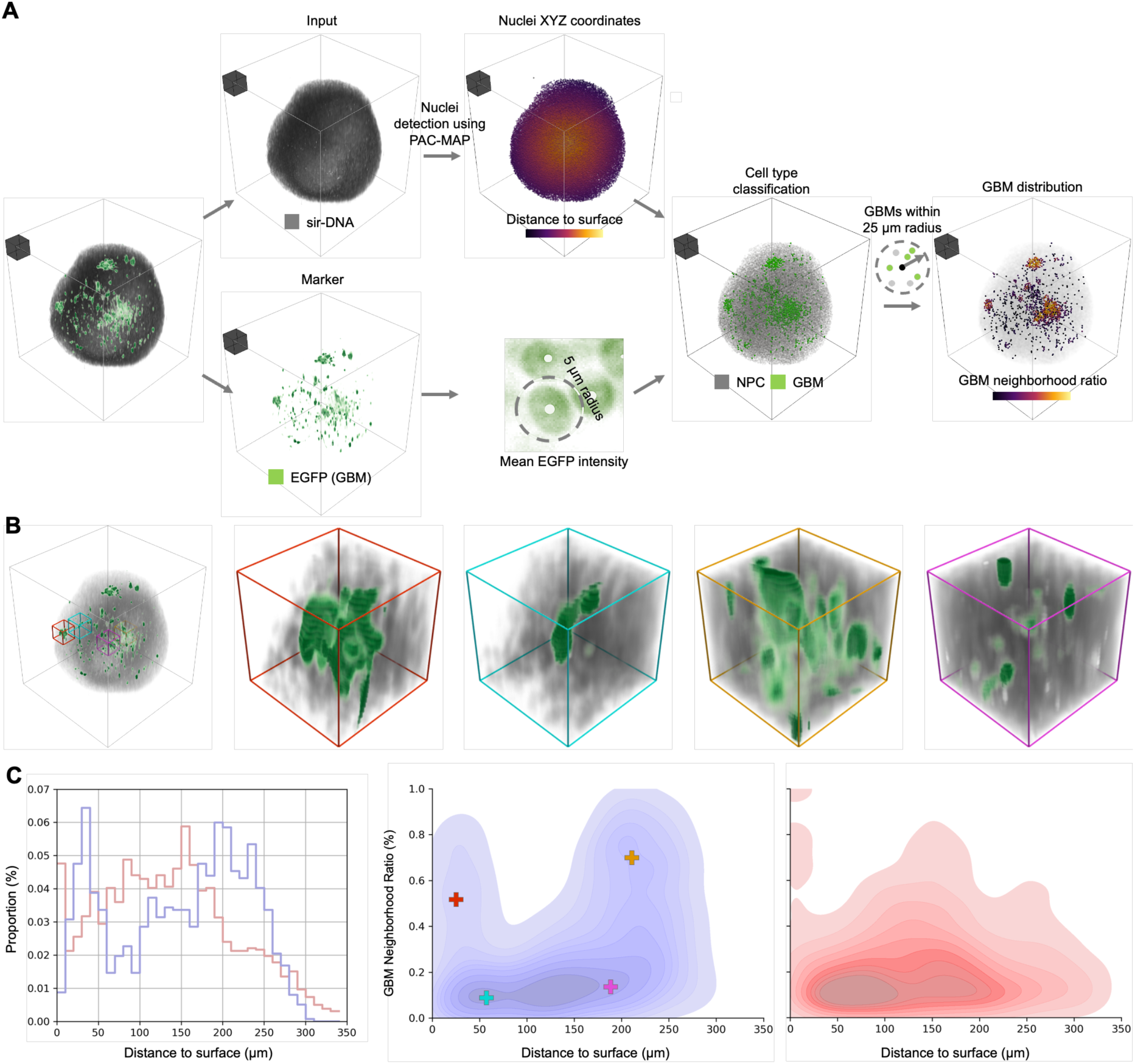
Mapping of GBM infiltration in cerebral organoid using instance centroid prediction. A: Image processing steps incuding nuclei centroid prediction, calculation of the distance to the surface, marker intensity measurement, cell type classification, and determination local GBM neighborhood ratio. A gray box of 100×100×100 μm^3^ is added to all renders as a scale reference; B: 3D renderings of a cerebral organoid infiltrated by GBM cells and subvolumes (100 × 100 × 100 µm^!^) from different radial positions and different local GBM concentration. The image was acquired 13 days after co-culturing 1000 GSCs with the organoid; C: GBM distribution visualized by histogram of GSCs and kernel density estimates of GBM neighborhood ratio as a function of distance to the surface. Blue: Organoid co-culture with 1000 GSCs. Red: Organoid co-cultured with 2000 GSCs. In the second kernel density plot, the median distance to the surface and GBM neighborhood ratio of the GSCs in the subvolumes in B are indicated.

## 4 Discussion

### 4.1 Instance centroid prediction is a robust and accurate strategy for nuclei detection

For 3D cell cultures, especially those with high cell density, the available instance segmentation methods often fail. However, many downstream tasks can also be achieved using positional information, *e.g.,* of nuclei centroids. To obtain such positional information, we developed a 3D U-Net to predict nuclei centroid probability maps in which centroids are identified by detecting local maxima (C-MAP). The main advantage of our method is that it is easy to implement on new datasets, as it only requires point annotations for training. We showed that this approach outperformed state-of-the-art cell detection strategies ([19,20]) and a pretrained 3D Stardist model on multiple datasets.

Our method differs from [19] in that we directly predict centroids instead of a binary foreground mask from which centroids are deduced, and that we train using relatively unfiltered masks (weak supervision) instead of eroded manually annotated masks to maximize performance in function of annotation effort. Similar to our approach, [20] proposed to use a 3D U-Net-based model to predict cell density maps for cell counting and detection. However, individual instances were identified by thresholding the predicted cell density map and, similar to [19], performing connected component analysis. Both [19] and [20] mention most false negatives were due to touching nuclei. Instead, we identify individual instances as peaks in the predicted centroid probability maps, which showed to improve recall up to +17 % in dense LSFM spheroid images.

### 4.2 Proximity adjusted centroid probability maps allow for adaptive local maxima finding

Additionally, we introduced the use of a proximity adjusted centroid probability map (PAC-MAP) that allows for adaptive local maxima finding, and which showed to be either on par or slightly improve performance (up to +1.3 % in F1) relative to C-MAP. PAC-MAP alleviates the need for having to set a fixed minimal allowed reference distance between peak candidates in the local maxima finding step. Since during training the appropriate local distance threshold is computed from the point annotations, and in inference it is predicted by the model, it removes the need for setting the hyperparameter and thus might be more convenient for applying it to new datasets compared to PAC-MAP. The proximity adjusted probability map was inspired by the spatial embeddings that are used in training instance segmentation models [4,12], e.g., gradients or vector fields that allow for delineating individual cells via post-processing, but it has the benefit that it can be computed from point annotations.

### 4.3 Weak supervised pretraining is an effective solution for small-data regimes

PAC-MAP showed better performance on the mouse brain LSFM dataset than on the SH-SY5Y spheroid datasets. While we attribute most of the difference to the more challenging nature of the high cell density spheroid datasets, it should also be noted that there were significant differences in the number of annotated nuclei in the training data. The mouse brain training dataset had around 14k annotated nuclei, while the SH-SY5Y training datasets had less than 3K. With the SH-SY5Y spheroids, however, we showed that weakly supervised pretraining with a relatively large dataset of suboptimal targets from conventional image processing techniques, and then finetuning on a smaller dataset of manual annotations can significantly improve performance compared to training from scratch. From this, a workflow can be suggested in which model training is first bootstrapped to weak targets, *e.g.,* suboptimal predictions obtained from an available conventional image processing algorithm. Thereafter, the model can be iteratively finetuned using a human-in-the-loop approach [27] with limited ground truth training data obtained by manually correcting predictions of its previous state.

Remarkably, For PAC-MAP, the performance gains of pretraining (+3.0 %) were lower compared to C-MAP (+4.5 %). Potentially the difference is caused by the increased impact of errors in the weak target centroid probabilities when they are weighted by the proximity to the nearest neighbor. While for C-MAP an error in the weak targets remains restricted to the specific instance, in PAC-MAP an error affects the intensities of multiple instances simultaneously. This makes it more challenging for the model to discover the true signal in the weak targets and might result in less adapted model weight initialization for finetuning.

### 4.4 Limitations

We designed PAC-MAP with simplicity in mind. While PAC-MAP performs well, there might be ways to further improve its performance, which have not been tested here. One option could consist of scaling the centroid probabilities by the average distance to the ￼*k* (￼*k*, with *k* > 1. In pretraining, the average ￼*k*against false positives/negatives in the weak targets than using only the distance to the nearest neighbor. However, that might make the method less performant in samples with high cell density heterogeneity. Second, for simplicity we decided on jointly predicting centroid position and proximity to the nearest neighbors in a single output channel. In future work, it could be investigated if separating centroid probabilities and proximities to nearest neighbors in *k* + 1 output channels. Herein, one channel could be reserved for centroid probabilities, while the other *k* output channels would store the distances to the nearest neighbors in *k* fixed directions. While this approach would require a more advanced post-processing algorithm, it might improve centroid candidate selection, *i.e.,* reducing false positives, via a crowdsourced voting mechanism. However, since touching nuclei is the most challenging problem, in which the proximity to the nearest neighbor affects recall the most, we opted for our current approach. Next to these technical aspects, several application-specific challenges remain, such as the detection of dividing cells, binucleated cells (*e.g.,* cardiomyocytes, cancer cells) or cells with extreme nuclear atypia, which were not explored in this work. Depending on the required behavior (cell *vs* nuclei detection), manual annotation should be adjusted, or pooling of multiple nuclei into a single cell object via post-processing might be needed. The latter could be handled by another downstream model tailored to the specific application.

## 5 Conclusion

We developed PAC-MAP, a 3D U-net based method to accurately predict the position of nuclei centroids in 3D fluorescence microscopy images. It provides a solution for analyzing dense samples when available instance segmentation models fail, and insufficient training data is available for improving their performance. Our method is easy to implement on new datasets, as it only requires point annotations for training. We showed that the combination of pretraining with weak supervision by a conventional image processing-based approach and supervised finetuning with human annotations is an effective strategy to improve performance and limit the required amount of annotated data. Finally, we demonstrated proof-of-concepts for using PAC-MAP in quality control and composition analysis of spheroids and organoids. We believe PAC-MAP can form the basis for more generic in toto cell composition analysis of 3D cell systems, such as cell counting, cell phenotyping, cell-to-cell or cell-to-tissue proximity analysis and capturing tissue organization.

## Acknowledgements

We thank Dr. Jorrit De Waele (University of Antwerp, Belgium) and Prof. Frederik De Smet (KU Leuven, Belgium) who kindly provided us respectively the LN-18-RED cell line and GSCs used for generating the samples for the proof-of-concept experiments.

## Data availability

The original generated data underlying this article are available on Zenodo, at https://dx.doi.org/10.5281/zenodo.11636385.

The mouse brain LSFM datasets were derived from the public Bitbucket repository at https://bitbucket.org/steinlabunc/numorph/downloads/3DNucleiTracingData.zip.

## Code availability

The code is available on GitHub, at https://github.com/DeVosLab/PAC-MAP.

