## Supplements for "PAC-MAP: Proximity Adjusted Centroid Mapping for Accurate Detection of Nuclei in Dense 3D Cell Systems"

**Supplementary material**

*Cell culture*

All cells were cultured at 37°C and 5 % $CO_2$. SH-SY5Y neuroblastoma cells were maintained in DMEM-F12 + Glutamax (Gibco, 10565018) supplemented with 10% Fetal Bovine Serum (Gibco, 10500064). The LN18-RED glioblastoma cell line was obtained by lentiviral transduction of LN18 cells to stably express RFP as a pan-cellular marker and was maintained in DMEM (Gibco, 41965039) supplemented with 10% Fetal Bovine Serum. Patient-derived glioma stem cells (GSC) (LBT037-EGFP, KU Leuven, Belgium) were maintained on laminin-coated plates (Sigma-Aldrich, L2020) in medium consisting of NeuroCult (Stemcell technologies, 05751) supplemented with Antibiotic/Antimycotic (Thermofisher, 15240062), 5 µg/ml FGF2, 5 µg/ml EGF and 2 µg/ml heparin (Stemcell technologies, 07980). Stable expression of EGFP by the GSCs was obtained via lentiviral transduction with a pLV[shRNA]-EGFP:T2A:Puro-U6-Scramble viral vector with EGFP construct and selection with puromycine (1µg/ml). The cell lines were subcultured upon 90% confluency using trypsin-EDTA (Gibco, 25200072). Human iPSCs (Sigma Aldrich, iPSC0028 Epithelial-1) were cultured on Matrigel (Corning, 734-1440) in Essential 8 medium (Gibco, A1517001). Upon cell thawing, 10µM Rock inhibitor (Y-27632 dichloride, MedChem, HY-10583) was added. Subculturing of iPSCs was performed with ReLeSR (Stemcell Technologies, 05872). Differentiation of iPSCs to neural progenitor cells (NPCs) was initiated by subculturing the iPSCs single cell using Tryple Express Enzyme (Life technologies, 12605010) at a density of 10e4 cells/cm$^2$ in mTesR1 medium (Stemcell Technologies, 85850) and Rock inhibitor. The following day, differentiation to NPCs was started by dual SMAD inhibition in neural maintenance medium (1:1 Neurobasal (Life technologies, 21103049):DMEM-F12 + Glutamax (Gibco, 10565018), 0.5x Glutamax (Gibco, 35-050-061), 0.5 % MEM Non Essential Amino Acids Solution (Gibco, 11140050), 0.5 % Sodium Pyruvate (Gibco, 11360070), 50 µM 2-

Mercaptoethanol (Gibco, 31350010), 0.025 % Human Insulin Solution (Sigma Aldrich, I9278), 0.5x N2 (Gibco, 17502048), B27 (Gibco, 17504044), 50 U/ml Penicillin-Streptomycin (Gibco, 15140122) supplemented with 1 µM LDN-193189 (Miltenyi, 130-106-540), SB431542 (Tocris, 1614). Daily medium changes were performed for 11 consecutive days. Following neural induction, the cells were matured in neural maintenance medium until DIV30 before organoid seeding. NPCs were subcultured using Tryple Express Enzyme.

*Obtaining weak targets for pretraining via conventional image processing algorithm*

In the conventional image processing algorithm, the spheroid volumes were first binarized (foreground) using the triangle threshold algorithm followed by a 3D binary opening (kernel size = 9), hole filling, the removal of small objects (< 1000 voxels), and a 2D median filter (kernel size = 5, *i.e.,* majority voting in a binary image) in XY. Thereafter, the binary volumes were divided in patches as earlier described. Then, inside of the foreground mask of every patch, the nuclei were segmented by using the Niblack threshold algorithm on the corresponding grayscale patch. Next, a 3D binary opening and a 2D median filter in XY (kernel size = 1 µm) were performed to smoothen the nuclei mask. Thereafter, a distance map was generated using the Euclidian distance transform and a Gaussian filter (kernel size = 1 µm). Using the voxel size, the kernel sizes were expressed in voxels, and rounding upwards. Thereafter, local maxima were found in the distance map using scikit-image's peak_local_max algorithm that was adapted to work with anisotropic images. In brief, voxel coordinates of peak candidates were transformed to physical coordinates and candidates were filtered based on distance in physical space. Only peaks ≥7.5 µm away from the image border and from other peaks were retained. Finally, the peaks were used as seeds for the seeded watershed algorithm and the centroids of the segmented objects were used as nuclei centroid targets.
